# Aperiodic Exponent as an EEG Marker of Interoceptive Attention during Gastric Fullness

**DOI:** 10.64898/2026.11.09.765525

**Authors:** Sabrina Liu, Nandini Datta, James D. Lock, Todd P. Coleman

## Abstract

Gastric interoceptive attention to internal, bodily signals such as hunger, satiety, and fullness are important for regulating feeding behaviors. However, abnormal processing or sensitivity to these can lead to disorders that result in overly restrictive or excessive consumption, highlighting the need for better understanding of the cortical processing of these cues. Specifically, understanding cortical representations of self-perceived gastric fullness have yet to be studied in humans. In this work, we demonstrate that consistently across individuals, the broadband power spectral density of EEG signals, as quantified by the aperiodic exponent, flattens after individuals drink water until fullness. Simultaneous measures of the stomach and heart demonstrate increased vagal drive to peripheral organs. We further find that changes in cortical aperiodic exponent patterns are associated with changes in the frequency of gastric oscillations. Overall, this suggests that the EEG aperiodic exponent may be a useful noninvasive biomarker for assessing gastric interoceptive attention.

## 1 Introduction

Gastric sensations play an important role in managing intake of food and water. These interoceptive cues can provide information about internal nutrient, caloric, and hydration levels that are used systemically across the body. Beyond that, gastric sensations can also provide mechanosensitive information about the gastrointestinal system itself and can be modulated by the volume of gastric distension and the rate of digestion. Clinically, processing interoceptive cues is important for studying patients with eating disorders (EDs), where aberrant interoceptive processing may contribute to or result from these conditions [1]. In other contexts, circulating glucagon-like peptide-1 (GLP-1) is important in sensing glucose levels in the blood and triggering satiation signals in the brain [2] while peripherally delaying gastric emptying and slowing gut motility [3]. The recent popularity of GLP-1 receptor agonists has generated interest in better understanding how these pharmaceuticals impact interoceptive processes involving satiation cues [4]. Thus, decoding multi-organ interactions between the brain and peripheral organs are pivotal to comprehensively understanding physiological responses to and the generation of gastric interoceptive cues.

Prior work has implicated parieto-occipital regions in processing gastric sensations. For example, in resting state, the phase of alpha waves in this region have been shown to be coupled to the amplitude of the gastric slow wave [5], and a similar region has been shown to present event-related potentials in response to vibratory pills in the stomach [6].

One challenge in understanding neural signatures of gastric interoception—and interoception in general—is being able to deliver and measure controlled stimuli to peripheral organs. Beyond the vibratory pill used in [6], many studies have used balloon distension to probe the impact of stomach distension [7, 8]. Previous studies have shown that the extent of gastric distension can lead to different neural signatures. Specifically, [8] uses fMRI data to demonstrate that only at high volumes of balloon distension was there additional activation of the left posterior amygdala. This region is a part of the limbic system and has been shown to be associated with emotional processing of food.

Though these devices allow more nuanced investigative control, they present gastric stimuli under non-naturalistic conditions that may not occur during self-regulated eating and drinking behavior. They can thereby introduce confounds to probing interoception, such as introducing pain, hyper-localized distension in the stomach, and distension rates that are abnormally fast or slow. These additional sensations may generate physiological responses that are different from natural consumption. In addition, from a practical experimental design perspective, when working with different patient groups, these tools may require additional analgesics or the consumption of a device that may be more difficult to tolerate for certain groups or age categories. For literature that uses water loading, most ask participants to consume a fixed volume [9] or drink until satiety [10]. These may generate sensations that do not draw as much conscious attention compared to a more salient state such as fullness.

Gastric fullness provides an opportunity to study interoceptive cues when they are more salient in one’s consciousness than in other states. This state may lead to heightened interoceptive attention towards internal bodily signals [11]. Traditionally, the alpha power in electroencephalography (EEG) has been used as a marker of attention [12]. Recently, there has been increased interest in another neural feature that may be a useful biomarker in this context: the aperiodic exponent. Prior work has studied the aperiodic exponent during a heartbeat counting task, which probes cardiac interoception [13]. They showed that the aperiodic exponent drops more when individuals participate in an exteroceptive auditory task than when attention is focused internally on heartbeat counting. In the context of respiratory interoception, it has been shown that the aperiodic exponent is modulated depending on whether or not breathing is volitional or spontaneous [14]. Together, this motivates studying the aperiodic exponent, in addition to oscillatory features such as alpha power that have been implicated in gastric coupling and attention, in the context of gastric fullness.

**Fig. 1.**
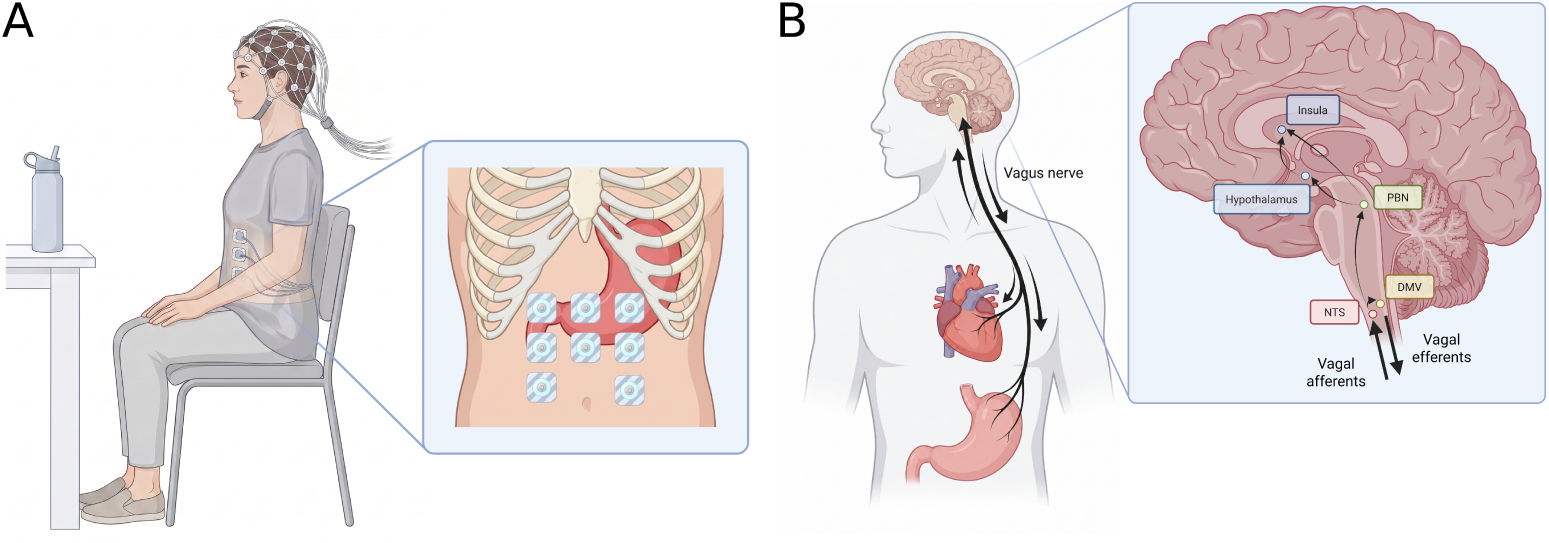
A) Experimental setup. Participants are fitted with a 128-channel EEG net to measure cortical signals. In addition, they have 8 recording electrodes on their stomach for EGG from which we can extract and measure gastric and cardiac signals. Participants are seated, eyes open during the experiment. For water loading, they drink from a straw from an opaque water bottle to minimize exteroceptive cues of water consumption. B) We propose modulation of peripheral and cortical signals is driven in part by a vagovagal response to water loading. Created in BioRender. L, S. (2027). https://BioRender.com/112es7c and https://BioRender.com/m6kse6z

In this work, we studied central and peripheral electrophysiologic changes in 34 participants aged 8 to 22 years old. Participants performed a two-stage water loading task (WLT-II), where they were first asked to drink to satiation and then drink to self-perceived fullness [15]. We recorded simultaneous resting state electroencephalography (EEG) and electrogastrography (EGG) for 15 minutes before and after water loading. We studied changes in the EEG aperiodic exponent as well as in features of the gastric slow wave and cardiac timings to characterize the physiological response to fullness in both the brain and peripheral organs. After water loading (i.e., drinking) to fullness, we demonstrated that consistently across individuals, the aperiodic exponent decreases. Simultaneous measures of the stomach and heart demonstrate increased vagal drive to the stomach (as evidenced by an increase in gastric oscillatory power and a decrease in gastric oscillatory frequency) and to the heart (as evidenced by increases in heart rate variability and other autonomic cardiac measures). We further found that changes in EEG aperiodic exponent was associated with changes in the frequency of gastric oscillations.

## 2 Results

### 2.1 Study overview

34 participants aged 8-22 years (17.96 *±* 4.16 years) were recruited to perform a two-stage water loading task (WLT-II) [15]. In this task, they drank water first until satiation. Then, they drank until they reached self-perceived fullness. The volumes consumed in each stage were recorded. For this task, participants were fitted with 128-channel EEG and 8-recording channel EGG. They were recorded at resting state before the task for 15 minutes. After reaching fullness, they were recorded for another 15 minutes. EGG analyses used data from the upper left (relative to the participant) electrode, which is expected to be closest anatomically to the stomach [16].

### 2.2 Aperiodic exponent decreases after drinking water

To assess if there were any cortical differences between the fasted and full states, we extracted the aperiodic exponent using the “Fitting Oscillations and One Over F” or FOOOF [17] procedure. We applied FOOOF to the pre- and post- recording separately for each participant on each of the 128 EEG electrodes. We then computed the average aperiodic exponent across the scalp and showed the changes in Fig. 2A. There was on average a *−*0.0913 decrease in the aperiodic exponent after drinking water. This was deemed significant by a one-sided, paired t-test (*p* = 0.00576).

**Fig. 2.**
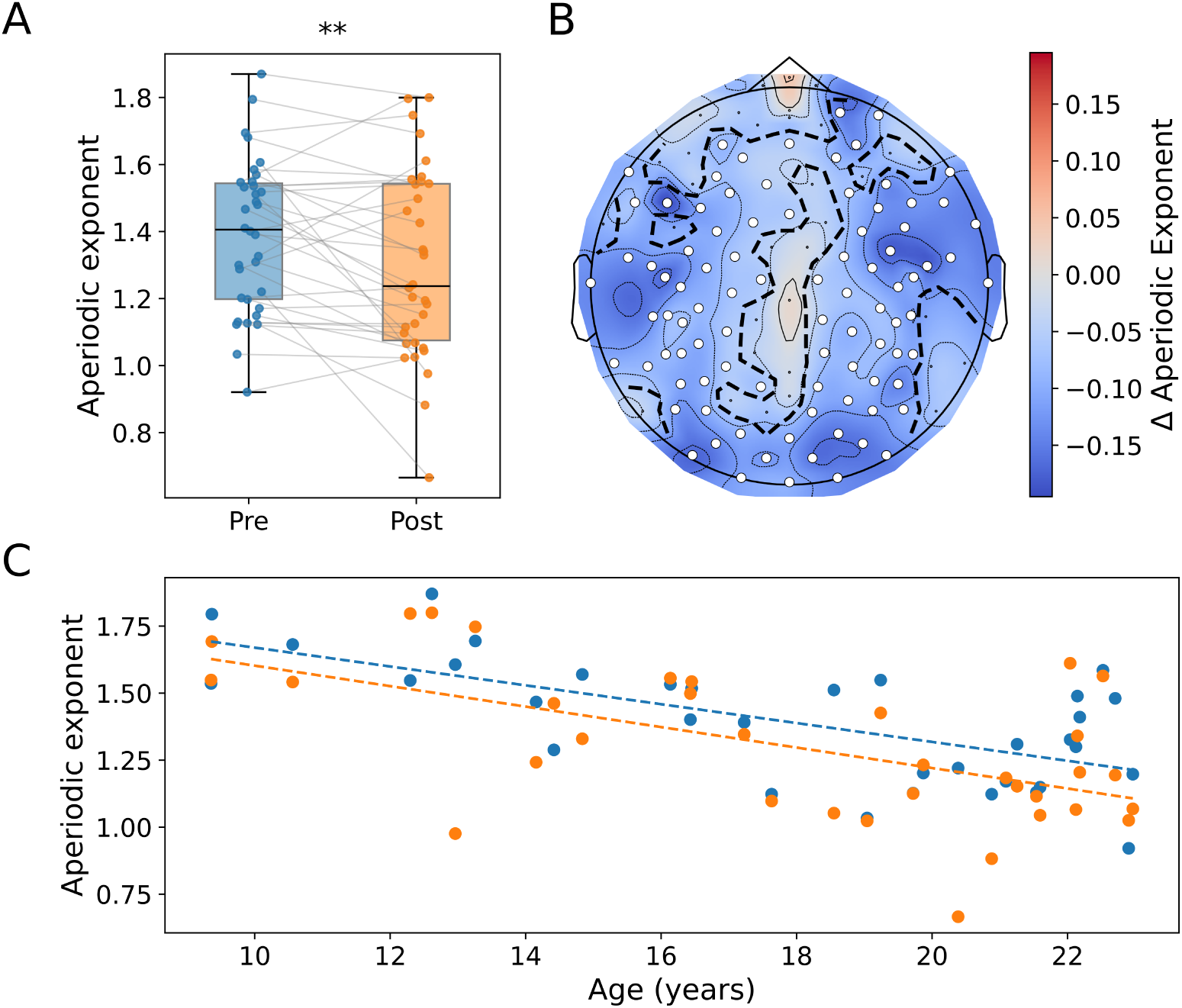
A) There is a significant drop in the aperiodic exponent after drinking water (*p* = 0.00576, one-sided, paired t-test). B) Topoplot showing the average per electrode change after drinking water. A significant cluster, as determined by the cluster-based permutation test, is outlined by the black dashed lines and indicated by white circles (*p* = 0.0036). C) Linear regression showing the effects of age on the aperiodic exponent in the pre- (blue) and post-drink (orange).

To identify which groups of electrodes contribute to these changes in the aperiodic exponent, we used nonparametric cluster-based permutation testing procedures [18]. As seen in Fig. 2B, one cluster widely distributed across the scalp exhibited a significant aperiodic exponent decrease (*p* = 0.0036).

Prior work has demonstrated that the aperiodic exponent decreases with age [19–21]. Given the wide age range of our participants, we next determined whether these changes in the aperiodic exponent are affected by age. To do this, we fit a linear mixed effects model (LMEM) described by:

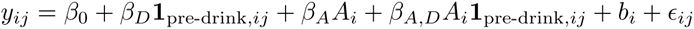

where *i* indexed the participant, while *j* indexed whether the sample was from the pre- or post-drink stage. *A_i_* was the participant’s age in years, and **1**_pre-drink*,ij*_ was an indicator for the drink stage. The coefficients *β*_0_, *β_D_*, *β_A_*, and *β_A,D_* captured the fixed effects associated with the intercept, the drink stage, the age, and the interaction between the two, respectively. *b_i_* captured participant-level random effects, while *ɛ_ij_* captured the residual error for each sample. The coefficients and statistics associated with this model are presented in Table 1. We also visualized the linear regression of the effect of age on the scalp-wide average aperiodic exponent pre- and post-drink in Fig. 2C. We showed that there was a significant decrease in aperiodic exponent with age. After accounting for age, there was also a significant decrease in the aperiodic exponent after water loading. The interaction between these two, i.e., does the amount of decrease change with age, did not significantly change.

**Table 1.** Coefficients and statistics from a linear mixed effects model testing the effects of age and drink stage on the aperiodic exponent.

| Effect | *β* | SE | p-value | 95% CI Low | 95% CI High |
|---|---|---|---|---|---|
| Intercept | 1.390 | 0.0352 | < 1 × 10<sup>−6</sup> | 1.321 | 1.459 |
| Age | −0.0352 | 0.00846 | 3.2 × 10<sup>−5</sup> | −0.0518 | −0.0186 |
| Drink | −0.0913 | 0.0346 | 0.00829 | −0.159 | −0.0235 |
| Age × drink | −0.00302 | 0.00830 | 0.716 | −0.0193 | 0.0133 |

### 2.3 Apparent changes in alpha power may be due to changes in the aperiodic component

Previous work has shown that there are changes in the alpha band that are associated with gastric interoceptive input [9]. Many of these did not explicitly account for the aperiodic component. Therefore, we tested whether there were alpha power changes even after considering the aperiodic component. To do that, we divided the 128 EEG electrodes into 5 common EEG regions shown in Fig. 3A. We used the alpha range defined as limits [8, 12]Hz. We considered three different methods for computing alpha power as depicted in Fig. 3B. First, we considered the uncorrected alpha power which takes the area under the power spectral density (PSD) within the specified band. It is also common to compute a normalized alpha power which takes the uncorrected value and divides that by the total area under the entire PSD. This normalized power adapts for changes in background power which can be caused by factors such as impedance changes, especially during longer recording sessions. Finally, we considered an aperiodic-adjusted alpha power that removes the aperiodic component and takes the area in the band of interest in what remains.

**Fig. 3.**
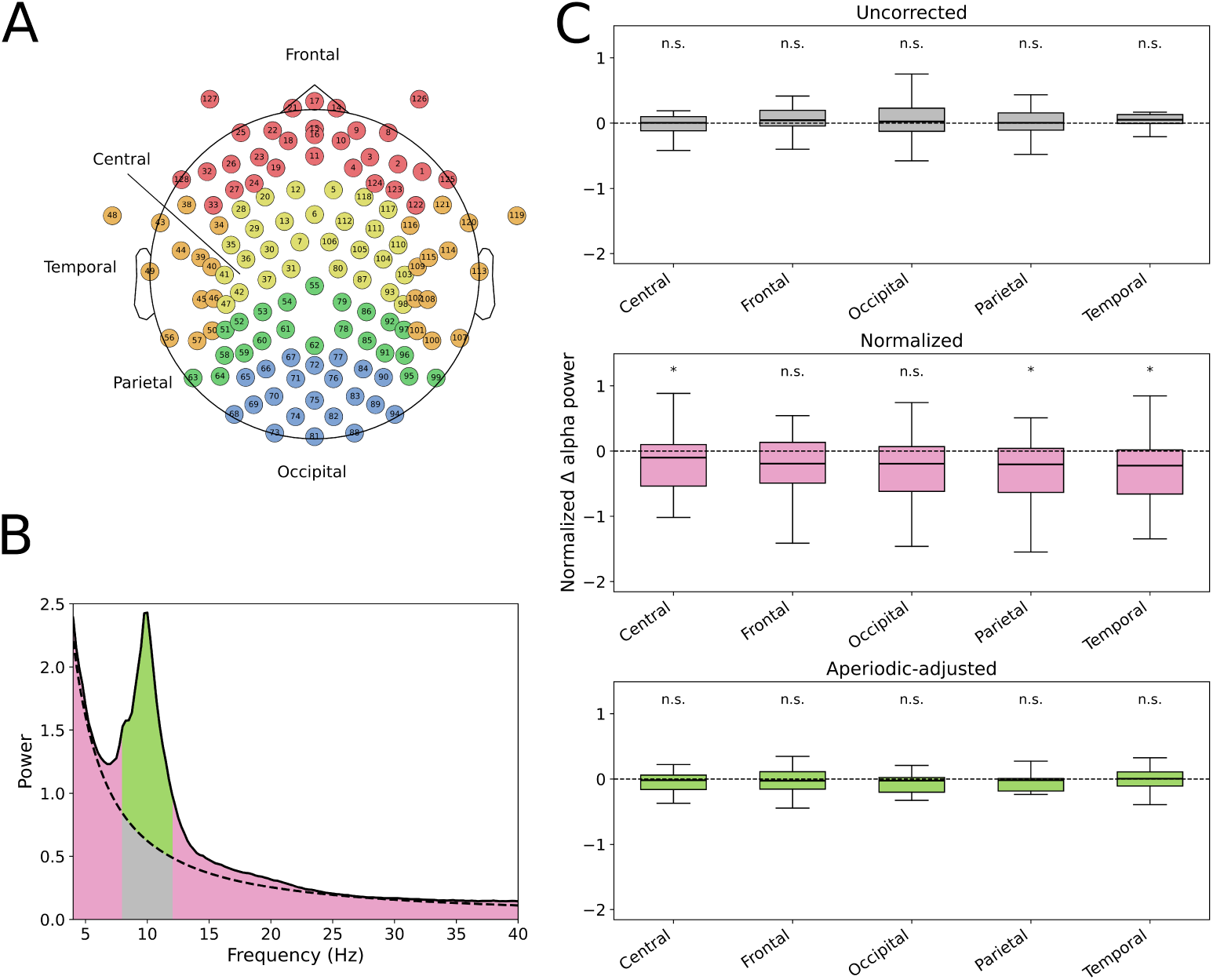
A) Layout of electrodes by regions, with frontal in red, temporal in orange, central in yellow, parietal in green, occipital in blue. B) Demonstrative figure for how the different alpha powers are computed. Uncorrected alpha power takes the area under the PSD (solid line) in the 8 to 12 Hz range, while normalized alpha divides that by the total area under the PSD. Adjusting for the aperiodic component would only take the area between the 8 to 12 Hz that is between the aperiodic fit (dashed) and the PSD (solid). C) The change in alpha power in each lobe between pre- and post-drink. From top to bottom, boxplots are shown for when the value is uncorrected, normalized by the power in the entire spectra, and when we account for the aperiodic component. The results from one-sided, paired t-tests corrected with BH show that when using the normalized alpha power, significant differences are identified in lobes that are otherwise not apparent after adjusting for the aperiodic component.

Fig. 3C shows box plots that depict the pre- to post- changes in each lobe depending on the method used to compute the alpha power. The values have been standardized to show the relative change, and p-values from one-sided, paired t-tests are used to determine significance after controlling the false discovery rate using BH across all the box plots shown. We showed that in none of the lobes do we have evidence of significant changes in the uncorrected alpha power. However, when normalized relative to the total power, the central, parietal, and temporal regions showed significant decreases. The aperiodic-adjusted alpha power, however, showed no evidence of significant changes in any lobes. This suggests that without properly correcting for the aperiodic component, there may be apparent narrowband power changes that are not necessarily due to changes in oscillatory activity alone.

### 2.4 Change in EGG dominant frequency is correlated with the water volume consumed

The human stomach has a slow wave frequency near 0.05 Hz. To see if there were gastric changes associated with water loading to fullness, we used the continuous wavelet transform to estimate the EGG dominant frequency. We also computed the normalized EGG power by taking the power within [0.03, 0.07]Hz and dividing that by the total power at that time from [0.01, 0.1]Hz [22].

In Fig. 4A, we averaged the dominant frequency and normalized power over time for each participant’s recording. After controlling the false discovery rate with Benjamini-Hochberg (BH) correction, we showed that there was a significant decrease in EGG frequency (one-sided, paired t-test, *p^∗^* = 0.00367) that was on average 0.001 46 Hz. In addition, there was also a significant increase in normalized EGG power (one-sided, paired t-test, *p^∗^* = 0.00367) on average of 0.0498.

**Fig. 4.**
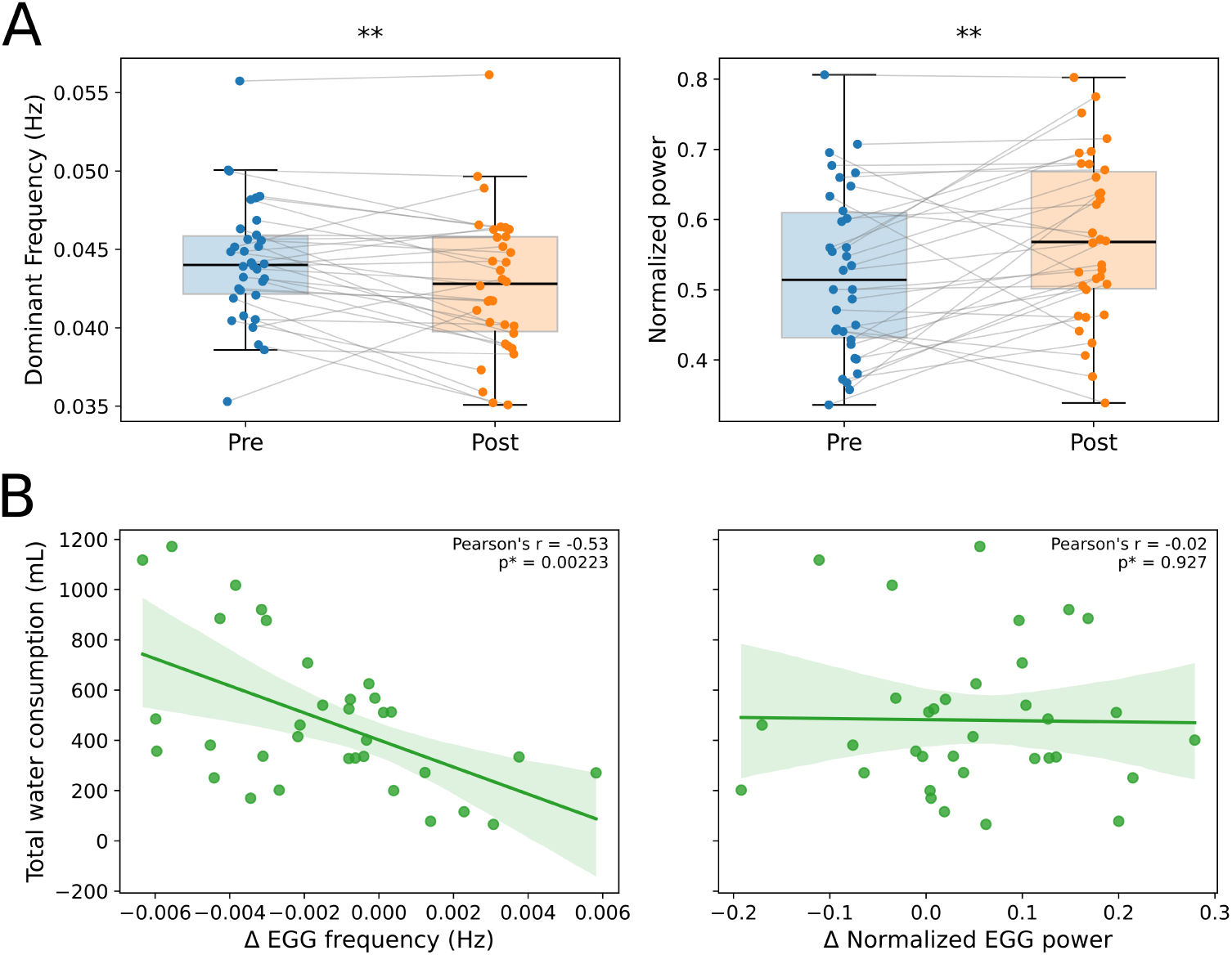
A) From left to right, change in EGG dominant frequency and normalized EGG power before and after drinking water. EGG dominant frequency significantly decreases (one-sided, paired t-test *p^∗^* = 0.00367) while EGG normalized power significantly increases (one-sided, paired t-test *p^∗^* = 0.00367). B) Correlations between EGG features with consumed water volume. There is a significant negative linear correlation between the consumed water volume with the change in EGG frequency (Pearson’s *r* = *−*0.53, *p^∗^* = 0.00223) while there is no significant correlation with the change in normalized EGG power (Pearson’s *r* = *−*0.02, *p^∗^* = 0.927).

When we correlated these with the total volume of water each individual consumed until fullness, we saw that the frequency drop was inversely correlated with total water consumed (Pearson’s *r* = *−*0.53, *p^∗^* = 0.00223). There was no significant linear correlation between water consumed and the change in normalized EGG power (Pearson’s *r* = *−*0.02, *p^∗^* = 0.927).

### 2.5 Increased parasympathetic activity to the heart after water loading

To also understand what changes in the peripheral nervous systems occur in response to water loading, from the EGG, we got the cardiac heartbeat timings by extracting the R peaks. From there, we computed the mean heart rate as well as quantified the heart rate variability (HRV) by computing the standard deviation in interbeat intervals (SDNN). We further computed an estimate of respiration rate using ECG-derived respiration [23].

In Fig. 5A, using one-sided, paired t-tests and after BH correction, we demonstrated a significant decrease in heart rate (*p^∗^ <* 0.001), a significant increase in HRV (*p^∗^ <* 0.001), and no evidence of significant change in respiration rate (*p^∗^* = 0.78).

**Fig. 5.**
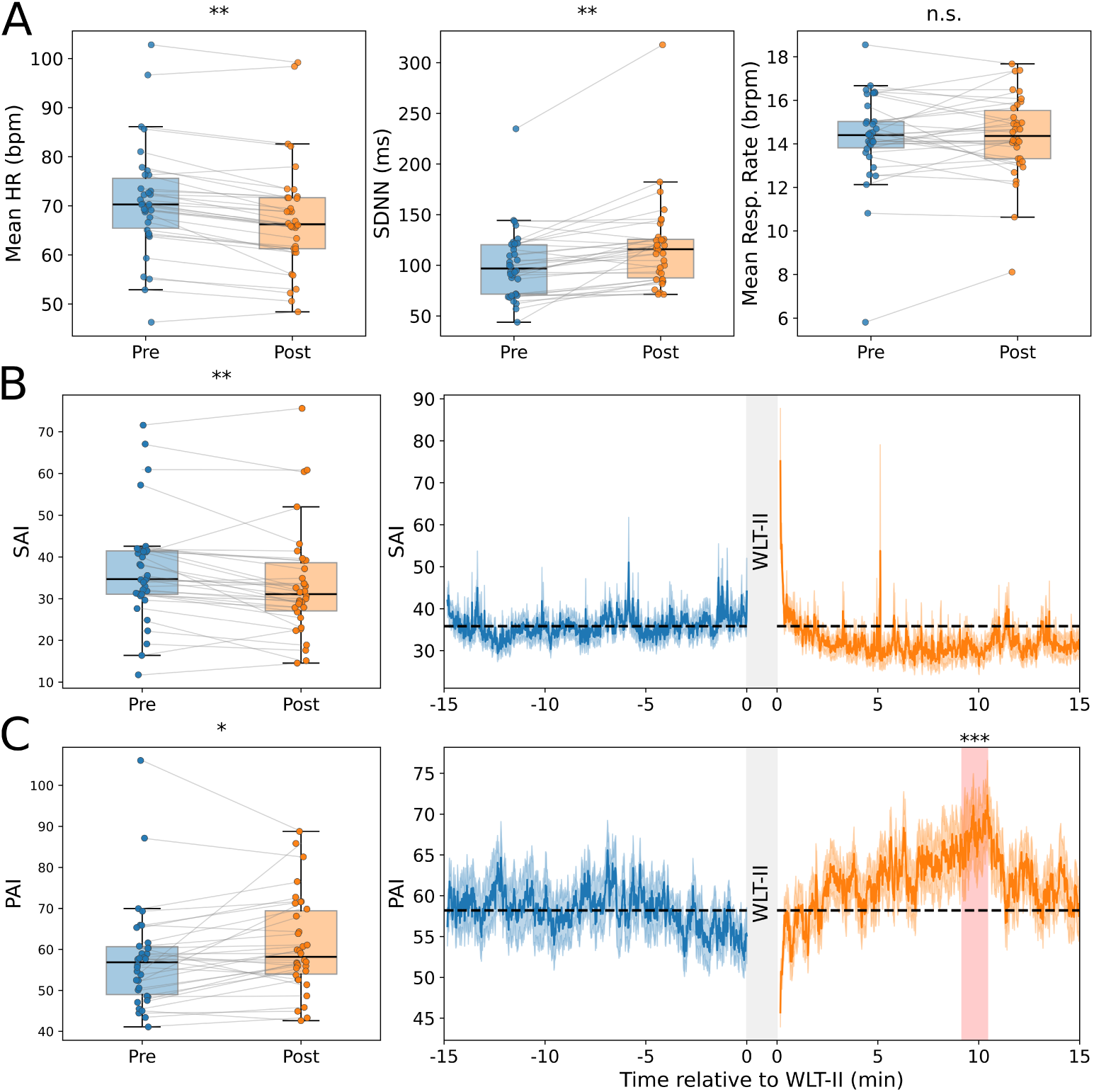
A) Changes in A) mean heart rate (*p^∗^* = 4.66 *×* 10*^−^*^8^), heart rate variability quantified by SDNN (*p^∗^* = 9.40 *×* 10*^−^*^5^), and respiration rate (*p^∗^* = 0.78) after water loading with significance determined by one-sided, paired t-tests corrected with BH. B) and C) show the the change in SAI (*p^∗^* = 0.0011) and PAI (*p^∗^* = 0.011), respectively, after drinking water. Time-resolved changes in SAI and PAI are shown. The mean is shown by the solid line with shaded bands indicating *±*1 SEM. The pre-drink mean is shown by the black, dashed line. One-sided cluster-based permutation testing identified a period of significant increase in PAI from baseline as highlighted in red from 9.17 to 10.43 minutes (*p* = 0.0004).

These changes are consistent with increases in parasympathetic innervation to the heart. We further computed more direct measures of autonomic innervation to the heart using the sympathetic activity index (SAI) and parasympathetic activity index (PAI) based on the extracted RR intervals [24]. In Fig. 5B and C, we showed that there was a statistically significant decrease in sympathetic activity (*p^∗^* = 0.0011) and a statistically significant increase in parasympathetic activity after water loading to fullness (*p^∗^* = 0.011). Dynamic measures of cardiac autonomic innervation, as shown in Fig. 5, indicated a significant increase in PAI spanning around 9.17 to 10.43 minutes after drinking to fullness as determined by cluster-based permutation testing (*p* = 0.0004).

### 2.6 Cortical aperiodic dynamics correlate with stomach response

To better understand how cortical aperiodic dynamics change over time and see if there are any patterns in how these changes occur throughout the scalp, we used Spectral Parameterization Resolved in Time (SPRiNT) to obtain a time-resolved estimate of the aperiodic exponent in each of the 128 electrodes for the pre- and post-drink stages [25]. We then performed principal component analysis (PCA) for each subject to obtain the top three components that accounted for the most variability in the combined pre- and post-drink data. Fig. 6A shows principal component trajectories from an example subject.

**Fig. 6.**
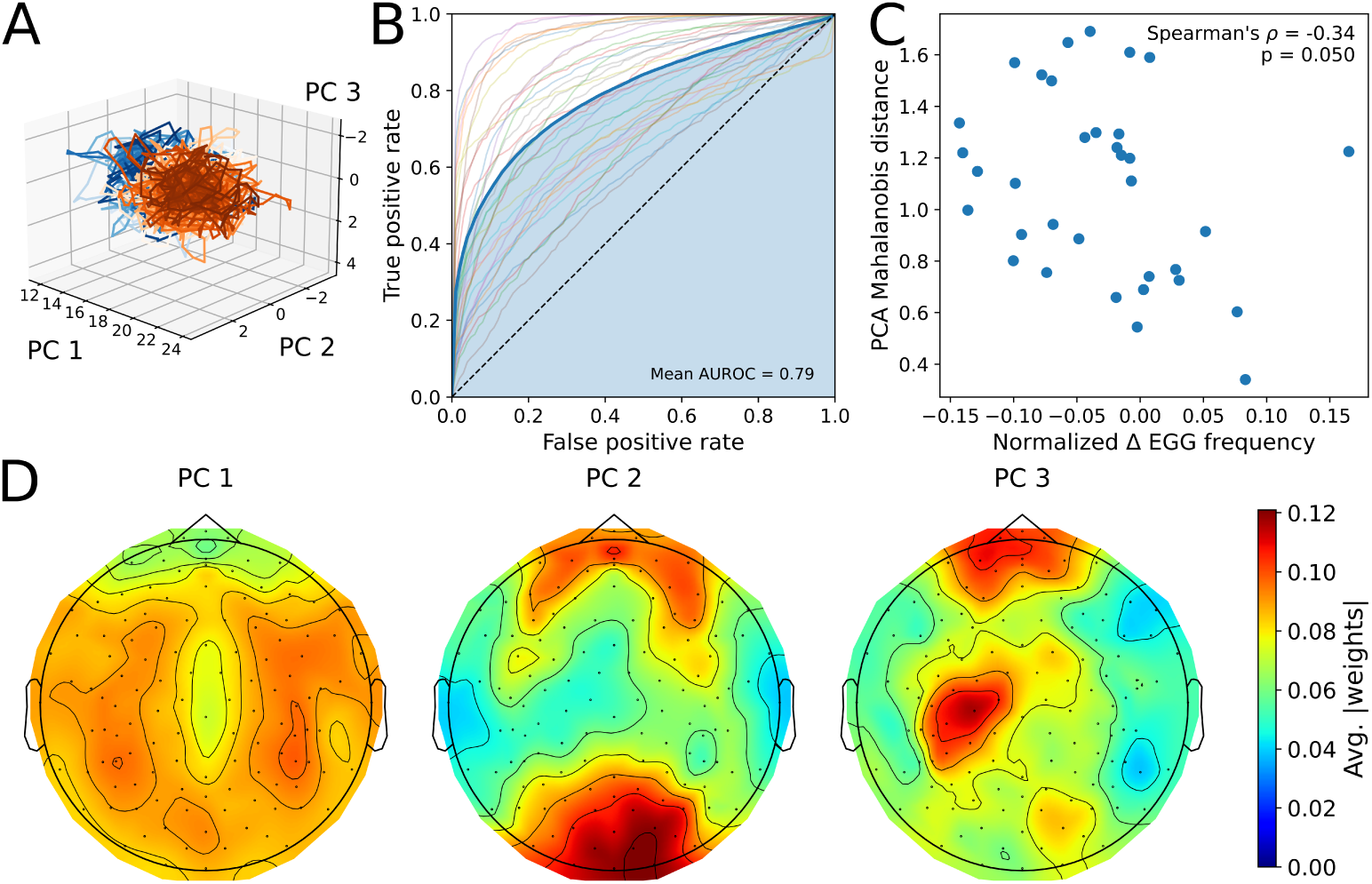
A) Example PCA trajectories from pre- (blue) and post- (orange) drink. B) AUC curves from logistic regression to classify drink stage based on the top 3 components with the average curve across all subjects shown in bolded blue. The mean AUROC was 0.79. C) There was a moderate negative association between normalized change in EGG frequency and the Mahalanobis distance between the pre- and post-drink trajectories (Spearman’s *ρ* = *−*0.34*, p* = 0.050). D) Heatmaps of the average magnitudes of the PC weights for the top 3 components.

To determine if these time trajectories are able to provide information about whether the data is from pre- or post-drink at the individual subject level, we trained a logistic regression model on a subset of their pre- and post-drink recordings and then used it to classify what drink stages the held-out test data points were from. We performed 5-fold cross validation and showed in Fig. 6B that there was an average area under the receiver operating curve (AUROC) of 0.79. This suggests that, at the individual level, the top 3 PCs associated with cortical aperiodic dynamics provided information that distinguishes between pre- and post-drink.

Since there was variability between subjects on the separability of their pre- and post-drink trajectories, we computed the Mahalonobis distance to quantify how separable they were [26]. The greater the distance, the more distinct the pre- and post-drink trajectories were. When we correlated these distances with the gastric changes, we demonstrated that in Fig. 6C that there was a moderate negative association between this distance and the normalized change in EGG dominant frequency (Spearman’s *ρ* = *−*0.34*, p* = 0.050). This suggests that for participants with greater cortical aperiodic differences across drink stages, there was also a greater decrease in their gastric dominant frequency.

To visualize which scalp regions contribute the most to PCs correlated with gastric activity, we plotted the absolute weights of the top 3 components averaged across all the subjects in Fig. 6D. PC 1 was associated with widespread changes in the aperiodic exponent across the scalp that were similar to the changes seen in Fig. 2B. PC 2, however, received most of its contributions from the parieto-occipital region and frontal region. PC 3 received most of its contributions from the left central and frontal regions.

## 3 Discussion

We demonstrated that the EEG aperiodic exponent significantly decreased after water loading to fullness. The temporal dynamics of these changes revealed that they had a modest association with changes in the EGG dominant frequency. This suggests that the EEG aperiodic exponent may represent part of a cortical response to gastric interoceptive stimuli.

This widespread reduction in the aperiodic exponent after water loading to fullness may reflect increased cortical arousal. Previous work comparing different stages of sleep and wakefulness has shown that increased arousal is associated with a flatter power spectral density and reduced aperiodic exponent [27]. In addition, reductions in the aperiodic exponent have also been shown to occur in response to exteroceptive stimuli, including auditory and visual cues [28, 29]. However, for interoceptive stimuli, findings have been less clear. The aperiodic slope has been shown to change in response to different types of respiration [30]. However, a previous water loading study did not find a robust aperiodic change after drinking to satiety [10]. One potential explanation for the difference is the end state participants were asked to drink to. The more pronounced gastric sensations that come from fullness as opposed to satiety may bring greater interoceptive attention to internal bodily signals, possibly leading to the reduction in the aperiodic exponent.

Our work also highlights the importance of separately considering aperiodic and oscillatory components of EEG data. We showed regional differences in normalized alpha power were no longer statistically significant after adjusting for the aperiodic exponent, suggesting that the changes may be partially due to broadband differences. This motivates revisiting prior work that has shown decreased alpha power after participants consumed a fixed volume of water [9] and investigating whether such changes are due to contributions from the oscillatory activity, aperiodic component, or both. In addition, there has also been prior work demonstrating coupling between the gastric phase and alpha amplitude [31], prompting whether a similar relationship exists for the aperiodic component as well. Such work would expand our understanding of how gastric rhythms may be linked to cortical activity.

We found significant changes in the EGG dominant frequency and normalized power in response to water loading that were consistent with findings in previous studies [32]. We demonstrated that between these two features, the total volume of water consumed until fullness was more closely correlated with the EGG dominant frequency than the EGG normalized power. This may be due to additional factors such as the geometry of the stomach during distension that affects surface power measurements. Simultaneous invasive manometry and cutaneous EGG in humans have demonstrated that the increase in power is not due to increased stomach contractions [32]. It has been proposed that these power changes are instead at least partially due to distension of the stomach bringing it closer to the surface electrodes. The same work also demonstrated a reduction in EGG frequency in humans after water loading, and it has been shown in canines that there is a negative correlation between frequency reduction and the volume consumed [33]. The exact mechanisms for this slowing of the gastric slow wave are unclear. Experiments in isolated interstitial cells of Cajal (ICC) show that mechanical stretch alone leads to increased firing rates in these pacemaker cells [34], conflicting with the slowing that occurs after water loading. Inhibitory vagal pathways may play a role. While often associated with cholinergic, excitatory processes, there are also vagal efferents onto the stomach that are inhibitory and attach to enteric neurons expressing NOS1 [35]. It has been shown that the release of nitric oxide (NO) can slow the frequency of slow waves in the antrum in canines [36] and in the small intestine of rodents [37]. However, whether such a pathway may be involved in the gastric response to water loading in humans, and whether osmotic or mechanical triggers initiate such a process, remains to be studied.

The temporal dynamics of the aperiodic exponent also provided potential evidence connecting cortical and gastric responses to water loading. After dimensionality reduction using PCA, there was a moderate association between the separability of the pre- and post-drink EEG recordings and the relative change in EGG dominant frequency. This suggests these cross-organ changes may reflect related processes. The first component had weights that were widespread throughout the scalp. The second component, however, demonstrated stronger contributions from parieto-occipital regions, which are also implicated in stomach-brain coupling at resting state [5] and event-related potentials in response to localized gastric stimuli [6]. All in all, these observations are consistent with afferent signaling from the stomach, though we were unable to eliminate potential causes due to other effects of water loading.

Together, our multimodal recordings suggest that there are gastric, cardiac, and cortical changes after water loading to fullness. Prior work establishes a vagovagal pathway that outlines a potential mechanism to describe such changes [35]. Specifically, gastric sensory information travels along afferent fibers in the vagus nerve to the nucleus tractus solitarius (NTS) while signals for autonomic regulation of peripheral organs descend along efferent fibers [35, 38]. The observed cardiac changes are consistent with vagally mediated parasympathetic input, and these mechanisms can also lead to inhibitory vagal efferents that slows the gastric slow wave frequency. At the same time, the reductions in the aperiodic exponent seen broadly across the scalp are consistent with increased cortical arousal. This suggests that parasympathetic, “rest- and-digest” peripheral patterns can coexist with heightened processing in the brain as attention is directed towards salient gastric sensations. However, our recordings could not directly test these pathways or probe if these changes were due to interoceptive attention alone. Future work comparing responses to different levels of gastric cues, like satiety vs. fullness, could help determine whether the aperiodic exponent can be a useful tool in distinguishing these states.

There are several limitations to our study. While water loading provides a relatively noninvasive method to distend the stomach, it does not isolate the mechanical stimulus. Gastric osmoreceptors have been shown to detect different solute concentrations in the stomach and regulate gastric emptying rates [39]. In addition, drinking water leads to a pressor response that engages the sympathetic nervous system, initiated by the renal nerve and sustained by the splanchnic nerve [40, 41]. Part of the parasympathetic changes may reflect compensatory mechanisms to regulate the cardiovascular system in response. Therefore, we cannot disentangle mechanical and osmotic effects that may contribute to the observed cortical and peripheral changes. Further work with varying fluid composition could help distinguish these effects.

Fullness in our study was through subjective self-reports. In addition, participants were permitted to drink for as much time as they wanted to reach this state. While this allowed for more individual specificity, it also introduced more variability in the time duration between the start and end of water loading. Different gastric emptying during this period could lead to differences in physiological responses. Additionally, without an additional method to objectively assess gastric distension, we were unable to verify if all participants reached comparable states of fullness. Using imaging such as with ultrasound could help clarify how subjective fullness, distension, and the physiological changes are related.

This study demonstrates that the aperiodic exponent of the EEG may be a potential indicator of cortical response to gastric interoceptive stimuli. We show that water loading to fullness is associated with scalp-wide changes in the aperiodic exponent, and these changes are associated with changes in the EGG dominant frequency. This highlights the importance of future work to analyze the aperiodic component along-side traditional oscillatory measures, thereby characterizing responses to interoceptive stimuli. Comparing measures of satiety and fullness across clinical populations, including individuals with EDs who may experience altered interoceptive processing, stands to clarify how well each measure captures subjective gastric sensations and how these relationships vary with interoceptive functioning.

## 4 Methods

### 4.1 Experimental Protocol

34 participants aged 8-22 years old were recruited for the study as approved by the Institutional Review Board at Stanford University. The participants were required to have no history of or any current psychiatric diagnoses or gastrointestinal conditions. In addition, they were required to be able to read and understand English. Participants that were 18 years or older provided informed consent. Those that were younger than 18 years old provided assent while their parents or legal guardians provided informed consent. Study data were collected and managed using REDCap electronic data capture tools hosted at Stanford University [42, 43]. Details about the average participant demographics are provided in Table 2.

**Table 2.** Summary demographics of participants. Shown are the mean and standard deviation values.

|  | Demographics Summary (*N* = 34) |
|---|---|
| Age | 17.96 ± 4.16 yrs |
| Male: Female | 11 : 23 |
| Height | 64.09 ± 4.54 in |
| Weight | 124.51 ± 29.27 lbs |
| Body Mass Index (BMI) | 20.30 ± 4.55 |

Each participant was asked to fast, including no water, for at least 8 hours before the study visit. All visits were held in the morning to coincide with an overnight fast. Participants were instrumented with an 8-channel OpenBCI data acquisition system which was placed on the abdomen to extract the EGG signal [22, 44, 45] as well as EGI’s 128-channel Geodesics EEG system at the Stanford University Koret Human Neurosciences Community Laboratory. 15 minutes of pre-drink resting state data were recorded where the participant was asked to sit upright and still with eyes open while looking straight in front of them. During this time, any additional auditory or visual stimuli in the recording room were minimized to reduce exteroceptive input.

After, participants were asked to perform the two-stage water load task (WLT-II), which requires them to first drink until satiation, or the state associated with termination of drinking and satisfaction with how much was consumed, and then to fullness [15]. Participants were given as much time as they needed to reach self-determined fullness. However, we prevented any external cues of consumed water volume by using opaque bottles placed on a table in front of the participant. The participant was not allowed to lift the bottle to determine the weight of the water consumed and could only drink through a straw. Once they deemed they were full, 15 minutes of post-drink resting state data were recorded.

### 4.2 EEG Preprocessing

The EEG data was preprocessed using a pipeline developed in EEGLab [46]. The data was recorded at 500 Hz and then downsampled to 250 Hz before being average re-referenced. Peripheral channels that were not over brain areas (e.g., below the ears) tended to have a lot of movement artifacts and were removed. The data was notch filtered to remove power line noise at 60 Hz and then bandpassed to [1, 100]Hz.

To remove large motion artifacts, artifact subspace reconstruction (ASR) was applied [47]. For mid-length recordings that are longer than seconds-scale, event-related analyses but shorter than hours-scale, sleep data, rejecting all segments with motion artifacts can lead to excessive data loss. ASR has been proposed as a practical solution to overcome this challenge [48].

After that, through visual inspection, additional channels were excluded if they had markedly high or low power relative to the others. No more than 5 additional channels were excluded in each recording in this step. Independent component analysis (ICA) was applied, and components associated with ocular and cardiac artifacts as well as those that represented large baseline drifts were removed. These were chosen manually with guidance by ICLabel, an automated classifier trained on manual annotations [49]. No more than 6 components were excluded for each recording. After ICA, the data was average re-referenced, and channels that had been removed were interpolated using spherical splines [50].

### 4.3 Computing the EEG Spectral and Aperiodic Features

The FOOOF algorithm (version 2.0.0rc0) was used to extract the aperiodic and oscillatory components from the preprocessed EEG [17]. Power spectra were parameterized across the frequency range [4, 40]Hz. The peak width limits were set to be between 0.5 Hz and 12 Hz. The minimum peak height was set to 0 and no limit was placed on the maximum number of peaks that could be identified. FOOOF was applied to each EEG electrode, and the exponent corresponding to the aperiodic component was computed.

We considered three different methods to estimate alpha power in EEG recordings. First, the uncorrected alpha power was computed by taking the area under the PSD in the range 8 Hz to 12 Hz. Second, normalized frequency bands were also employed, where the uncorrected power is then divided by the total frequency under the PSD. This helped correct for any broadband changes in power that could result from factors such as impedance changes during longer EEG recordings. Finally, we considered an aperiodic-adjusted alpha power. This approach takes the aperiodic fit extracted from FOOOF and subtracts it from the PSD. The total area under the curve in this difference was used.

When applicable, lobes were assigned by taking the EGI 128 channel Hydrocel GSN projection to a 10-10 system and extending that to include all electrodes [51].

### 4.4 EGG Analysis

We recorded the electrogastrogram (EGG) signal using an OpenBCI Cyton board. The data was collected at a sampling rate of 250 Hz and had 8 channels. The stomach is located underneath the participant’s left ribcage. For this study, we used one channel of EGG and selected the top left channel, since in most participants, it exhibits the highest gastric power [16].

The stomach slow wave oscillates at a frequency near 0.05 Hz, though frequencies ranging from 0.03 Hz to 0.07 Hz are considered normogastric. The EGG data was preprocessed by first detrending it by subtracting a linear least squares fit. Then, a Wiener filter was applied for artifact rejection [22]. The size of the window was set to the number of samples in a 0.03 Hz wave to avoid subtracting out the longest possible normal gastric signal. The signal was then decimated to 5 Hz.

To estimate the dominant EGG frequency at each time point, we employed the continuous wavelet transform (CWT) using a complex Morlet wavelet [52]. Wavelet powers at each time and frequency were computed by taking the squared magnitudes of the CWT coefficients. Naively taking the frequency with the greatest power at each time point will potentially extract frequencies that are outside of physiologically feasible ranges for the stomach. In addition, it can lead to sharp jumps in the frequency estimate from one time point to the next that would also not be physiologically feasible. To circumvent these issues, we developed a dynamic programming-based method which used a cost function that took into account these physiological constraints. Similar frameworks for ridge selection in time-frequency estimates have been developed in prior work [53]. We first identified all the local maxima in wavelet power. The cost function preferred frequencies near 0.05 Hz and that corresponded to these local maxima. Candidate paths were restricted to be within 0.02 Hz to 0.075 Hz with maximum frequency jumps of 0.005 Hz between time points. Dominant frequency estimates at each time point were taken from the path that had the minimum cost while obeying these constraints.

To estimate the power change in EGG due to water ingestion, we used the normalized EGG power [22], which took the power in the range 0.03 Hz to 0.07 Hz and divided it by the total power in the PSD.

### 4.5 Cardiorespiratory Analysis

From the raw EGG recordings before filtering or downsampling, R peaks were present in the waveforms, which allowed us to compute cardiac measures that are based on interbeat timings [45]. We used the Pan-Tompkins algorithm to identify the R peaks [54]. In time segments where the algorithm failed to identify all the peaks due to motion artifacts and other noise, we manually corrected the timings.

From these interbeat intervals, we computed the mean heart rate across each pre- and post-recording for each individual. In addition, we computed the standard deviation of the interbeat intervals (SDNN) across each recording as a measure of heart rate variability. While we did not utilize a respiration belt, we estimated respiration rate by using ECG-derived respiration (EDR) [23]. This algorithm uses the fact that chest movement from breathing can cause baseline wandering in the ECG waveform that can be extracted. We used the version that has been implemented in NeuroKit2 [55] to extract the breathing rate before and after water loading.

In addition to these methods, we obtained more direct measures of autonomic innervation to the heart using the sympathetic and parasympathetic activity indices (SAI and PAI) based upon point process statistical models of heartbeat dynamics [24]. Traditional spectral methods that estimate autonomic activity by looking at the low and high frequency bands of RR intervals have been shown to not always reliably separate sympathetic and parasympathetic activity [56]. SAI and PAI have been shown under a variety of autonomic stimuli to more accurately track these changes [24]. We computed the SAI and PAI timeseries in the 15 minutes pre-drink and the 15 minutes post-drink. To check for time periods with significant changes compared to baseline, we performed cluster-based permutation testing in the post-drink period relative to the mean of the pre-drink.

### 4.6 Time-Resolved Aperiodic Exponent

For each subject, we also extracted a time-resolved estimate of the aperiodic exponent. This was computed using the Spectral Parameterization Resolved in Time (SPRiNT) procedure [25]. This procedure applies FOOOF to PSDs computed using the short-term Fourier transform (STFT) in short windows of data. The FOOOF parameters used in each window were the same as the ones that were applied to the entire recording. As suggested in [25] when applying SPRiNT to longer datasets, each STFT was computed using 5 windows that were 2 s long with 75% overlap. In addition, to prune outlier peaks, those that had less than 3 peaks that were within 2.5 Hz in the 6 nearby time bins were removed.

Based on these time-resolved aperiodic exponent estimates for each electrode, we then applied principal component analysis (PCA) to reduce the dimensions from the 128 dimensional timeseries and choose the 3 components that accounted for the most variability [57, 58]. To do this, we applied the procedure to the combined data from pre- and post-drink for each subject individually. This created a shared basis between the stages for each subject and allowed us to see if there were components that can be used to separate the two.

To determine if these components carry information that distinguishes between pre- and post-drink states, for each subject, we performed 5-fold cross validation by training a logistic regression model for each subject. We computed the area under the receiver operator characteristic curve (AUROC) to quantify the accuracy.

In addition, we quantified how distinct the trajectories were by using the Mahalonobis distance between them [26]. This value measured the distance between the centers of the trajectories normalized by the variation in each of the trajectories. Those that were very distinct and separable should have distances between their centers that were large relative to the spread of each trajectory, increasing the Mahalanobis distance. This was compared to the normalized change in EGG dominant frequency. By normalizing, we were able to characterize how large the frequency change relative to the pre-drink frequency at baseline.

To visualize which EEG electrodes contribute the most to each of the principal components, we computed the PC weights from each subject for the top 3 components and plotted the average absolute value of these weights across all subjects to avoid canceling out positive and negative contributions.

## Acknowledgments

We would like to thank all of our participants and their families for being involved in the study. We would also like to thank Dr. Milena Kaestner and the staff at the Koret Human Neurosciences Community Lab for their technical support with the EEG recordings.

## Declarations

## References

[1] Khalsa, S.S., Berner, L.A., Anderson, L.M.: Gastrointestinal interoception in eating disorders: charting a new path. Current Psychiatry Reports 24(1), 47–60 (2022)

[2] Watts, A.G., Kanoski, S.E., Sanchez-Watts, G., Langhans, W.: The physiological control of eating: signals, neurons, and networks. Physiological Reviews 102(2), 689–813 (2022) 10.1152/physrev.00028.2020. PMID: 34486393

[3] Shah, M., Vella, A.: Effects of GLP-1 on appetite and weight. Reviews in Endocrine and Metabolic Disorders 15(3), 181–187 (2014)

[4] Kim, K.S., Park, J.S., Hwang, E., Park, M.J., Shin, H.Y., Lee, Y.H., Kim, K.M., Gautron, L., Godschall, E., Portillo, B., et al.: GLP-1 increases preingestive satiation via hypothalamic circuits in mice and humans. Science 385(6707), 438–446 (2024)

[5] Rebollo, I., Devauchelle, A.-D., Béranger, B., Tallon-Baudry, C.: Stomach-brain synchrony reveals a novel, delayed-connectivity resting-state network in humans. eLife 7, 33321 (2018)

[6] Mayeli, A., Al Zoubi, O., White, E.J., Chappelle, S., Kuplicki, R., Morton, A., Bruce, J., Smith, R., Feinstein, J.S., Bodurka, J., et al.: Parieto-occipital ERP indicators of gut mechanosensation in humans. Nature Communications 14(1), 3398 (2023)

[7] Hollerbach, S., Hudoba, P., Fitzpatrick, D., Hunt, R., Upton, A.R., Tougas, G.: Cortical evoked responses following esophageal balloon distension and electrical stimulation in healthy volunteers. Digestive Diseases and Sciences 43(11), 2558–2566 (1998)

[8] Wang, G.-J., Tomasi, D., Backus, W., Wang, R., Telang, F., Geliebter, A., Korner, J., Bauman, A., Fowler, J.S., Thanos, P.K., et al.: Gastric distention activates satiety circuitry in the human brain. Neuroimage 39(4), 1824–1831 (2008)

[9] Hallschmid, M., Mölle, M., Fischer, S., Born, J.: EEG synchronization upon reward in man. Clinical Neurophysiology 113(7), 1059–1065 (2002)

[10] Kalahasti, D.V., Sazonov, E., Malaia, E.: Identification of hunger and satiety states from EEG data. In: 2024 46th Annual International Conference of the IEEE Engineering in Medicine and Biology Society (EMBC), pp. 1–5 (2024). IEEE

[11] Joshi, V., Graziani, P., Del-Monte, J.: The role of interoceptive attention and appraisal in interoceptive regulation. Frontiers in Psychology 12, 714641 (2021)

[12] Ray, W.J., Cole, H.W.: EEG alpha activity reflects attentional demands, and beta activity reflects emotional and cognitive processes. Science 228(4700), 750–752 (1985)

[13] Fló, E., Belloli, L., Cabana, Á., Ruyant-Belabbas, A., Jodaitis, L., Valente, M., Rohaut, B., Naccache, L., Rosanova, M., Comanducci, A., et al.: Predicting attentional focus: Heartbeat-evoked responses and brain dynamics during interoceptive and exteroceptive processing. PNAS Nexus 3(12), 531 (2024)

[14] Karjalainen, S., Kujala, J., Parviainen, T.: Neural activity is modulated by spontaneous and volitionally controlled breathing. Biological Psychology 197, 109026 (2025)

[15] Van Dyck, Z., Vögele, C., Blechert, J., Lutz, A.P.C., Schulz, A., Herbert, B.M.: The Water Load Test as a Measure of Gastric Interoception: Development of a Two-Stage Protocol and Application to a Healthy Female Population. PloS one 11(9), 0163574 (2016)

[16] Riezzo, G., Russo, F., Indrio, F.: Electrogastrography in adults and children: the strength, pitfalls, and clinical significance of the cutaneous recording of the gastric electrical activity. BioMed Research International 2013(1), 282757 (2013)

[17] Donoghue, T., Haller, M., Peterson, E.J., Varma, P., Sebastian, P., Gao, R., Noto, T., Lara, A.H., Wallis, J.D., Knight, R.T., et al.: Parameterizing neural power spectra into periodic and aperiodic components. Nature Neuroscience 23(12), 1655–1665 (2020)

[18] Maris, E., Oostenveld, R.: Nonparametric statistical testing of EEG-and MEG-data. Journal of Neuroscience Methods 164(1), 177–190 (2007)

[19] Merkin, A., Sghirripa, S., Graetz, L., Smith, A.E., Hordacre, B., Harris, R., Pitcher, J., Semmler, J., Rogasch, N.C., Goldsworthy, M.: Do age-related differences in aperiodic neural activity explain differences in resting EEG alpha? Neurobiology of Aging 121, 78–87 (2023)

[20] McSweeney, M., Morales, S., Valadez, E.A., Buzzell, G.A., Yoder, L., Fifer, W.P., Pini, N., Shuffrey, L.C., Elliott, A.J., Isler, J.R., et al.: Age-related trends in aperiodic EEG activity and alpha oscillations during early-to middle-childhood. NeuroImage 269, 119925 (2023)

[21] Hill, A.T., Clark, G.M., Bigelow, F.J., Lum, J.A., Enticott, P.G.: Periodic and aperiodic neural activity displays age-dependent changes across early-to-middle childhood. Developmental Cognitive Neuroscience 54, 101076 (2022)

[22] Gharibans, A.A., Smarr, B.L., Kunkel, D.C., Kriegsfeld, L.J., Mousa, H.M., Coleman, T.P.: Artifact rejection methodology enables continuous, noninvasive measurement of gastric myoelectric activity in ambulatory subjects. Scientific Reports 8(1), 5019 (2018)

[23] Van Gent, P., Farah, H., Van Nes, N., Van Arem, B.: HeartPy: A novel heart rate algorithm for the analysis of noisy signals. Transportation Research Part F: Traffic Psychology and Behaviour 66, 368–378 (2019)

[24] Valenza, G., Citi, L., Saul, J.P., Barbieri, R.: Measures of sympathetic and parasympathetic autonomic outflow from heartbeat dynamics. Journal of Applied Physiology 125(1), 19–39 (2018)

[25] Wilson, L.E., Silva Castanheira, J., Baillet, S.: Time-resolved parameterization of aperiodic and periodic brain activity. eLife 11, 77348 (2022)

[26] McLachlan, G.J.: Mahalanobis distance. Resonance 4(6), 20–26 (1999)

[27] Lendner, J.D., Helfrich, R.F., Mander, B.A., Romundstad, L., Lin, J.J., Walker, M.P., Larsson, P.G., Knight, R.T.: An electrophysiological marker of arousal level in humans. eLife 9, 55092 (2020)

[28] Waschke, L., Donoghue, T., Fiedler, L., Smith, S., Garrett, D.D., Voytek, B., Obleser, J.: Modality-specific tracking of attention and sensory statistics in the human electrophysiological spectral exponent. eLife 10, 70068 (2021)

[29] Podvalny, E., Noy, N., Harel, M., Bickel, S., Chechik, G., Schroeder, C.E., Mehta, A.D., Tsodyks, M., Malach, R.: A unifying principle underlying the extracellular field potential spectral responses in the human cortex. Journal of Neurophysiology 114(1), 505–519 (2015)

[30] Kluger, D.S., Forster, C., Abbasi, O., Chalas, N., Villringer, A., Gross, J.: Modulatory dynamics of periodic and aperiodic activity in respiration-brain coupling. Nature Communications 14(1), 4699 (2023)

[31] Richter, C.G., Babo-Rebelo, M., Schwartz, D., Tallon-Baudry, C.: Phase-amplitude coupling at the organism level: The amplitude of spontaneous alpha rhythm fluctuations varies with the phase of the infra-slow gastric basal rhythm. NeuroImage 146, 951–958 (2017)

[32] Chen, J., McCallum, R.: Response of the electric activity in the human stomach to water and a solid meal. Medical and Biological Engineering and Computing 29(4), 351–357 (1991)

[33] Lin, H.C., Zhao, X.-T., Chung, B., Gu, Y., Elashoff, J.D.: Frequency of gastric pacesetter potential depends on volume and site of distension. American Journal of Physiology-Gastrointestinal and Liver Physiology 270(3), 470–475 (1996)

[34] Won, K.-J., Sanders, K.M., Ward, S.M.: Interstitial cells of cajal mediate mechanosensitive responses in the stomach. Proceedings of the National Academy of Sciences 102(41), 14913–14918 (2005)

[35] Berthoud, H.-R., Münzberg, H., Yu, S., Morrison, C.D., Kim, A., Neuhuber, W.L.: Anatomy and function of efferent and afferent vagal innervation of the stomach. Autonomic Neuroscience, 103465 (2026)

[36] Ozaki, H., Blondfield, D., Hori, M., Publicover, N., Kato, I., Sanders, K.: Spontaneous release of nitric oxide inhibits electrical, Ca2+ and mechanical transients in canine gastric smooth muscle. The Journal of Physiology 445(1), 231–247 (1992)

[37] Koh, S.D., Kim, T.W., Jun, J.Y., Glasgow, N.J., Ward, S.M., Sanders, K.M.: Regulation of pacemaker currents in interstitial cells of cajal from murine small intestine by cyclic nucleotides. The Journal of Physiology 527(1), 149–162 (2000)

[38] Levinthal, D.J., Strick, P.L.: Multiple areas of the cerebral cortex influence the stomach. Proceedings of the National Academy of Sciences 117(23), 13078–13083 (2020)

[39] Hunt, J.: Some properties of an alimentary osmoreceptor mechanism. The Journal of Physiology 132(2), 267 (1956)

[40] Routledge, H.C., Chowdhary, S., Coote, J.H., Townend, J.N.: Cardiac vagal response to water ingestion in normal human subjects. Clinical Science 103(2), 157–162 (2002)

[41] Mai, T.H., Garland, E.M., Diedrich, A., Robertson, D.: Hepatic and renal mechanisms underlying the osmopressor response. Autonomic Neuroscience 203, 58–66 (2017)

[42] Harris, P.A., Taylor, R., Thielke, R., Payne, J., Gonzalez, N., Conde, J.G.: Research electronic data capture (REDCap)—a metadata-driven methodology and workflow process for providing translational research informatics support. Journal of Biomedical Informatics 42(2), 377–381 (2009)

[43] Harris, P.A., Taylor, R., Minor, B.L., Elliott, V., Fernandez, M., O’Neal, L., McLeod, L., Delacqua, G., Delacqua, F., Kirby, J., et al.: The REDCap consortium: building an international community of software platform partners. Journal of Biomedical Informatics 95, 103208 (2019)

[44] Balasubramani, P.P., Walke, A., Grennan, G., Perley, A., Purpura, S., Ramanathan, D., Coleman, T.P., Mishra, J.: Simultaneous gut-brain electrophysiology shows cognition and satiety specific coupling. Sensors 22(23), 9242 (2022)

[45] Subramanian, S., Kunkel, D.C., Nguyen, L., Coleman, T.P.: Exploring the gut-brain connection in gastroparesis with autonomic and gastric myoelectric monitoring. IEEE Transactions on Biomedical Engineering 70(12), 3342–3353 (2023)

[46] Delorme, A., Makeig, S.: EEGLAB: an open source toolbox for analysis of single-trial EEG dynamics including independent component analysis. Journal of Neuroscience Methods 134(1), 9–21 (2004)

[47] Chang, C., Hsu, S., Pion-Tonachini, L., Jung, T.: Evaluation of artifact subspace reconstruction for automatic artifact components removal in multi-channel EEG recordings. IEEE Transactions on Biomedical Engineering 67(4), 1114–1121 (2019)

[48] Miyakoshi, M.: Artifact subspace reconstruction: a candidate for a dream solution for EEG studies, sleep or awake. Sleep 46(12) (2023)

[49] Pion-Tonachini, L., Kreutz-Delgado, K., Makeig, S.: ICLabel: An automated electroencephalographic independent component classifier, dataset, and website. NeuroImage 198, 181–197 (2019)

[50] Perrin, F., Pernier, J., Bertrand, O., Echallier, J.F.: Spherical splines for scalp potential and current density mapping. Electroencephalography and Clinical Neurophysiology 72(2), 184–187 (1989)

[51] Luu, P., Ferree, T.: Determination of the hydrocel geodesic sensor nets’ average electrode positions and their 10–10 international equivalents. Inc, Technical Note 1(11), 7 (2005)

[52] Komorowski, D., Pietraszek, S.: The Use of Continuous Wavelet Transform Based on the Fast Fourier Transform in the Analysis of Multi-channel Electrogastrography Recordings. Journal of Medical Systems 40(1), 10 (2016)

[53] Iatsenko, D., McClintock, P.V., Stefanovska, A.: Extraction of instantaneous frequencies from ridges in time–frequency representations of signals. Signal Processing 125, 290–303 (2016)

[54] Pan, J., Tompkins, W.J.: A real-time QRS detection algorithm. IEEE Transactions on Biomedical Engineering (3), 230–236 (1985)

[55] Makowski, D., Pham, T., Lau, Z.J., Brammer, J.C., Lespinasse, F., Pham, H., Schölzel, C., Chen, S.H.A.: NeuroKit2: A Python toolbox for neurophysiological signal processing. Behavior Research Methods 53(4), 1689–1696 (2021) 10.3758/s13428-020-01516-y

[56] Billman, G.E.: The LF/HF ratio does not accurately measure cardiac sympathovagal balance. Frontiers in Physiology 4, 26 (2013)

[57] Pearson, K.: LIII. On lines and planes of closest fit to systems of points in space. The London, Edinburgh, and Dublin Philosophical Magazine and Journal of Science 2(11), 559–572 (1901)

[58] Hotelling, H.: Analysis of a complex of statistical variables into principal components. Journal of Educational Psychology 24(6), 417 (1933)

